# Leveraging collaborations between researchers, wildlife managers and citizen-volunteers to monitor endangered carnivore populations across space and time

**DOI:** 10.64898/2026.09.16.752259

**Authors:** Arjun Srivathsa, Mayank Shukla, Pooja Saravanan, Divyajyoti Ganguly, Midhun Mohan, Uma Ramakrishnan

**Affiliations:** National Centre for Biological Sciences, TIFR, GKVK Campus, Bellary Road, Bengaluru 560065, India; Nature Conservation Foundation, Mysuru, Karnataka 570017, India

**Author notes:** Corresponding authors: Arjun Srivathsa; Uma Ramakrishnan, Arjun Srivathsa National Centre for Biological Sciences, TIFR, GKVK Campus, Bellary Road, Bengaluru 560065, India.

**Keywords:** collaborative monitoring, dhole, population density, single nucleotide polymorphisms, spatially explicit capture–recapture, Western Ghats

## Abstract

Monitoring populations is fundamental for conserving threatened carnivores, but can be difficult to sustain in the long term across large landscapes. We present a collaborative framework for monitoring dholes (*Cuon alpinus*) in India’s Western Ghats, combining fecal DNA-based individual identification, Spatially Explicit Capture–Recapture (SECR) models, state Forest Department participation, and trained citizen-volunteers. We surveyed across six sites between 2019–2023, encompassing three Protected Areas (Wayanad, Parambikulam, Periyar) and their adjacent Territorial Divisions. Fecal samples were genotyped using a panel of 96 Single Nucleotide Polymorphisms, and detections were analyzed using SECR models to estimate densities. In Wayanad Sanctuary, densities remained stable across 2019, 2022 and 2023, increasing marginally from 15.67 to 19.01 individuals per 100 sq.km. In 2023, densities varied substantially among six sites, from 9.73/100 sq.km in Parambikulam to 29.23 in Nemmara Territorial Division. Field implementation involved an average of 46 Forest Department personnel co-collecting data per site–year. Among former citizen-volunteers who were surveyed for project feedback (N=29), 83% went on to either work or pursue higher education in wildlife-related or-adjacent fields. We demonstrate that molecular individual identification coupled with SECR can provide a practical basis for dhole population assessments while enabling multi-stakeholder participation in field monitoring. Sustained engagement with wildlife managers and locally recruited volunteers can strengthen institutional capacity, facilitate knowledge exchange, and improve continuity of monitoring programs. We argue that integrating molecular ecology with collaborative governance and democratizing field-science offers a scalable pathway for landscape-level monitoring required to guide dhole conservation.

## 1. Introduction

Long-term and sustained monitoring has been acknowledged as being pivotal for understanding ecological systems, particularly for those that involve critical rare events, slow processes, high annual or periodic variability, or complex phenomena with multiple interacting dimensions (Franklin 1989; Magnuson 1990). Such monitoring programs, the world over, have greatly expanded human understanding of complex ecological relationships, population trends, multi-scale impacts of natural and human-induced stressors on populations, and detailed biological underpinnings of elusive species behavior––all of which are either difficult or near-impossible to observe in short-duration snapshot studies; this has also aided in formulating better-informed and evidence-based management mandates for species of conservation concern (Willis et al. 2007; Magurran et al. 2010; Huges et al. 2017). Two parallel realities become readily apparent when deliberating on long-term monitoring work: (i) these initiatives are able to capture high-quality information from large geographical areas, involve extensive human effort and resources, and generate massive amounts of extremely valuable data (e.g., COMADRE, Salguero-Gómez et al. 2016; also see Church et al. 2022); and (ii) the logistics, financial resources, infrastructure and human power required for implementing such work pose formidable challenges (Vucetich et al. 2020; Viblanc et al. 2026), especially in many countries of the Global South that struggle with limited resources and accessibility barriers in terms of institutional support and technical expertise (Rafiq et al. 2024; Scaini et al. 2024).

Scientists have long recognized the persistent and oftentimes wide gaps that exist between researchers studying ecological systems and wildlife management agencies that are tasked with protecting and managing them (Hulme 2014; Gosselin et al. 2018). Involving state agencies in biodiversity monitoring has its strengths: managers may possess valuable experiential knowledge that is frequently underutilized, anecdotal observations provide context for explaining ecological phenomena or generating testable hypotheses, and the diversity of actors allows for greater credibility and broadening *the ways of knowing* (Pouyat et al. 2010). Besides, co-produced knowledge is likely to have longevity and greater impact on management than externally driven research (Merkle et al. 2019; West et al. 2019). A sterling example, perhaps, is the United States Fish and Wildlife Service (USFWS), a federal entity tasked with studying *and* managing wildlands and wildlife. For a quarter of a century, USFWS has integrated rigorous scientific inquiry with adaptive management, together with periodic external reviews and audits to help rationalize resource allocation and optimize management outcomes (Ratz et al. 2005; Murphy and Weiland 2019). The adoption of such systems has been more recent in the Global South, where the most prolific model involves managers collecting field data (e.g., SMART patrols) while academic institutions or conservation NGOs provide backend analytical support (e.g., Kuiper et al. 2025; Sinchuri et al. 2025). Such examples hold promise, but are contrasted by ground realities in many places where wildlife bureaucracy–academic institution ties remain tenuous (see Velho et al. 2012).

The end of the 20th century heralded an age of increased public participation in ecological research by way of ‘citizen science’ (or alternatively, community-based science; Dickinson et al. 2012). Through these programs and initiatives, nature enthusiasts formed a new constituency of researcher-adjacent actors that helped exponentially scale-up biodiversity tracking and monitoring (Bonney et al. 2009; Aristeidou et al. 2021). Community-contributed studies were earlier limited to field-based co-collection of ecological data; subsequently, with the advent of and access to internet connectivity and smart devices, web-based elicitation of community-collected data became more mainstream (Sullivan et al. 2014; Callaghan and Gawlik 2015). Prominent examples from the world over include online platforms like iNaturalist and eBird, along with in-person initiatives like REEF (monitoring of marine fishes by volunteer SCUBA divers following structured training protocols); citizen-science projects have thus bridged several gaps between science and society, despite geographical biases titled towards North America, Europe and Oceania (Theobald et al. 2015). While the formative versions of this volunteering model largely entailed collection and curation of field observations, the past decade has also witnessed scientists expand the prospects to proactively and more intimately involve citizen-volunteers in research––like camera-trapping in their own backyards and neighborhoods (e.g., Laskey et al. 2021)––thereby *democratizing* field technology that was once exclusive to research institutions and wildlife agencies. Beyond offering opportunity for hands-on experience in research, volunteering for ecological monitoring has also helped create advocates for conservation among the general public, particularly for threatened species groups like large mammals (Forrester et al. 2017).

Mammalian large carnivores constitute a severely imperiled taxonomic group, with many species facing threats of extinction, globally (Fernández Sepúlveda and Martín 2022). Given their relatively long lifespans, peculiar life history traits and sensitivity to anthropogenic impacts, long-term studies of carnivores focused on demographic parameters like annual survival, recruitment and population growth rates are crucial from a conservation perspective (Balme et al. 2009; Chandler and Clark 2014; Sharma et al. 2014). Their rarity and global charismatic appeal bestow certain biases in favor of carnivores, in terms of funding, as well as research and conservation focus (Davies et al. 2018; Bellon 2019). Still, over decades, select carnivore species have disproportionately benefited from this kind of focus, while several others remain severely data-deficient (Brooke et al. 2014; Srivathsa et al. 2022; Strampelli et al. 2022). Studying large carnivores is also relatively restrictive in that it typically requires niche expertise, access to remote locations, specialized equipment and higher monetary resources (Boitani and Powell 2012). Because of these reasons, undertaking studies of carnivores has been exclusionary in certain cases, species, and locations (as compared to other taxonomic groups, like birds). The twin challenges of (i) restrictive access and (ii) many carnivores lacking baseline data on population vital rates can be potentially addressed by democratizing the science (e.g., Forrester et al. 2017) and implementing population monitoring exercises as multi-agency or cross-institutional undertakings (e.g., Van Der Weyde et al. 2021).

The Asiatic wild dog (‘dhole’ *Cuon alpinus*) is among the world’s most threatened large carnivores. Dholes are elusive, social canids that do not possess unique pelage patterns or other individually identifiable physical features. Most packs and populations inhabit dense forested landscapes across South and Southeast Asia. Estimating their populations has been a primary challenge for ascertaining their status across most parts of their range (Kamler et al. 2015; Srivathsa et al. 2020a, 2020b). While recent studies have explored various methodological approaches to estimate their numbers (Ngoprasert et al. 2019; Punjabi et al. 2022), standardized, robust, and replicable protocols for multi-year multi-site monitoring do not exist. We adopt the field, laboratory, and analytical methods proposed by Srivathsa et al. (2021) for fecal DNA-based individual identification of dholes and application of Spatially Explicit Capture–Recapture (SECR) models; we scale-up the approach to track dhole densities over time (3 years, one site) and space (6 sites, 1 year) in India’s Western Ghats. Rivas et al. (2026) outline five key components required for sustainable biodiversity monitoring–– (i) permanent infrastructure, (ii) local governance, (iii) integrated capacity building and training, (iv) global connectivity, and (v) diversified financing. Broadly following this framework, we leverage (i) efforts by researchers from a national academic institution, (ii) participation of state wildlife agencies, and (iii) training local citizen-volunteers, to show proof of concept for multi-stakeholder involvement in monitoring populations of an endangered large carnivore. We then discuss the ecological inferences and practical considerations, i.e., (iv) knowledge sharing and (v) funding models, relevant for sustaining such monitoring activities in the long term.

## 2. Materials and methods

### 2.1. Infrastructure and state agency participation

The project was conceived as a multi-year initiative (2019–2027) to estimate population sizes and model meta-population dynamics of dholes in India’s Western Ghats. Given that fecal DNA-based data were the only reliable source for individual identification of dholes, implementing the study hinged primarily on using molecular methods, necessitating institutional and equipment support that were conducive for the pertinent laboratory methods. The research was platformed at a national research institute in India with access to wet labs, in-house sequencing facilities and multiple high-performance computing clusters for bioinformatics analyses. The research was greenlit by the Kerala State Forest Department, with the Principal Chief Conservator of Forests/Chief Wildlife Warden (PCCF/CWW) providing research permits and mandating that the individual site-level department heads would serve as co-investigators for the field surveys. At the level of individual sites (Protected Areas and Territorial Divisions; detailed in the Study Area description below), the Divisional Forest Officers (DFOs) proactively facilitated the research through providing accommodation, provisioning food supplies (especially in remote locations with accessibility constraints), and assigning ground staff for carrying out field surveys with the research teams. The Forest Department ground staff––Forest Watchers, Tribal Watchers, Eco-Development Committee ‘EDC’ members, Nominal Muster Roll ‘NMR’ Watchers/Elephant Watchers, Beat Forest Officers (BFO) and Section Forest Officers (SFO) took part in the surveys and co-collected fecal DNA samples following field training prior to walking survey routes with researchers and citizen-volunteers. Here, we present a summary of the number of personnel involved annually, with a broad break down of their designations, from 2019 to 2025 (note that 2020 and 2021 had COVID-related lockdowns; no surveys were conducted during these two years).

### 2.2. Training and capacity building

Citizen-volunteer openings were advertised across social media and in-country job-based listservs as one–two-month paid internship positions (stipend: 5000INR/∼65USD per month, besides covering per diem food costs, accommodation and local travel). Applicants were shortlisted based on their academic background (biology/ecology/wildlife sciences), basic exposure to outdoor activities (experience in hiking, trekking, birdwatching or visits to Protected Areas), distance to the field site (preference given to applicants from Kerala, followed by neighboring states, to minimize travel costs) and proficiency in *Malayalam* (the local language of the state). Shortlisted applicants were contacted and briefed about the project, roles and responsibilities, field schedules, stipend, and logistical arrangements. The selected interns received comprehensive orientation covering the objectives of the project, field protocols, site maps, datasheets, sampling kits, and good field practices. Teams were also acquainted with Forest Department operations and organizational structure, prevention of sexual harassment (POSH) guidelines, and field safety practices. Field training included fecal sample collection, identification of indirect animal signs, navigation using hand-held GPS units, and respectful engagement with the department ground staff. Training and mentoring extended beyond the requirements of field surveys––interns were provided exposure to fundamentals of ecology and conservation through presentations, reading scientific literature and weekly paper discussions, the use of spreadsheets for data entry and validation, and software programs like QGIS for spatial data processing.

To assess the contribution of their internship toward their professional trajectories, 30 former interns who participated between 2019–2025 were sent feedback forms in June 2026 (ensuring a minimum of one-year duration since their tenure with the project). The questionnaire was provided in English and *Malayalam*. The questions pertained to the participants’ previous experience in wildlife-related projects/internships, current occupation, their perceptions on how this internship contributed to their understanding of scientific research and fieldwork, whether the stipend offered influenced their decision to participate, experience of working in a gender-balanced team, adequacy of the training provided prior to fieldwork, and whether their participation influenced their interest in pursuing higher education or a career in wildlife/wildlife-adjacent fields (**Supplementary File 1**). The survey questions did not include any personally identifiable information, and was implemented by a member of the project who was not involved with the current study.

### 2.3. Study area and field survey design

India’s Western Ghats landscape is a global biodiversity hotspot and a critical stronghold for dhole populations (Kamler et al. 2015; Srivathsa et al. 2020a, 2021; Rodrigues et al. 2022). We conducted field surveys in the southern part of this landscape, within the State of Kerala, across three Protected Areas (PAs): Wayanad Sanctuary (344 sq. km), Parambikulam (380 sq. km) and Periyar (925 sq. km), and three locations in the surrounding Territorial Divisions (TDs): Wayanad Territorial, Nemmara, and Kotttayam+Ranni with unprotected/multi-use reserve forests and agroforest plantations (**Fig. 1**). These PAs and the surrounding landscape mosaics are composed of evergreen and semi-evergreen forests, montane shola–grasslands, moist and mixed deciduous forests, and naturalized teak plantations, with some locations towards the lower elevations and northern latitudes also supporting dry deciduous forests with savannah woodlands; the commodity agroforests are dominated by tea and coffee, followed by smaller extents or patches of rubber and cardamom plantations (Das et al. 2006; Roy et al. 2015). All the forested areas (PAs and within TDs) are under the jurisdiction of the State Forest Department, while the coffee and tea agroforests are privately-owned plantations. The landscape supports high densities of dholes and their co-predators––tigers *Panthera tigris* and leopards *Panthera pardus* (see Jhala et al. 2020; Srivathsa et al. 2021), along with a suite of mid-to large-sized herbivore ungulates (gaur *Bos gaurus*, sambar *Rusa unicolor*, chital *Axis axis*, muntjac *Muntiacus muntjac* and wild boar *Sus scrofa*), which form the primary prey species for dholes.

**Figure 1.**
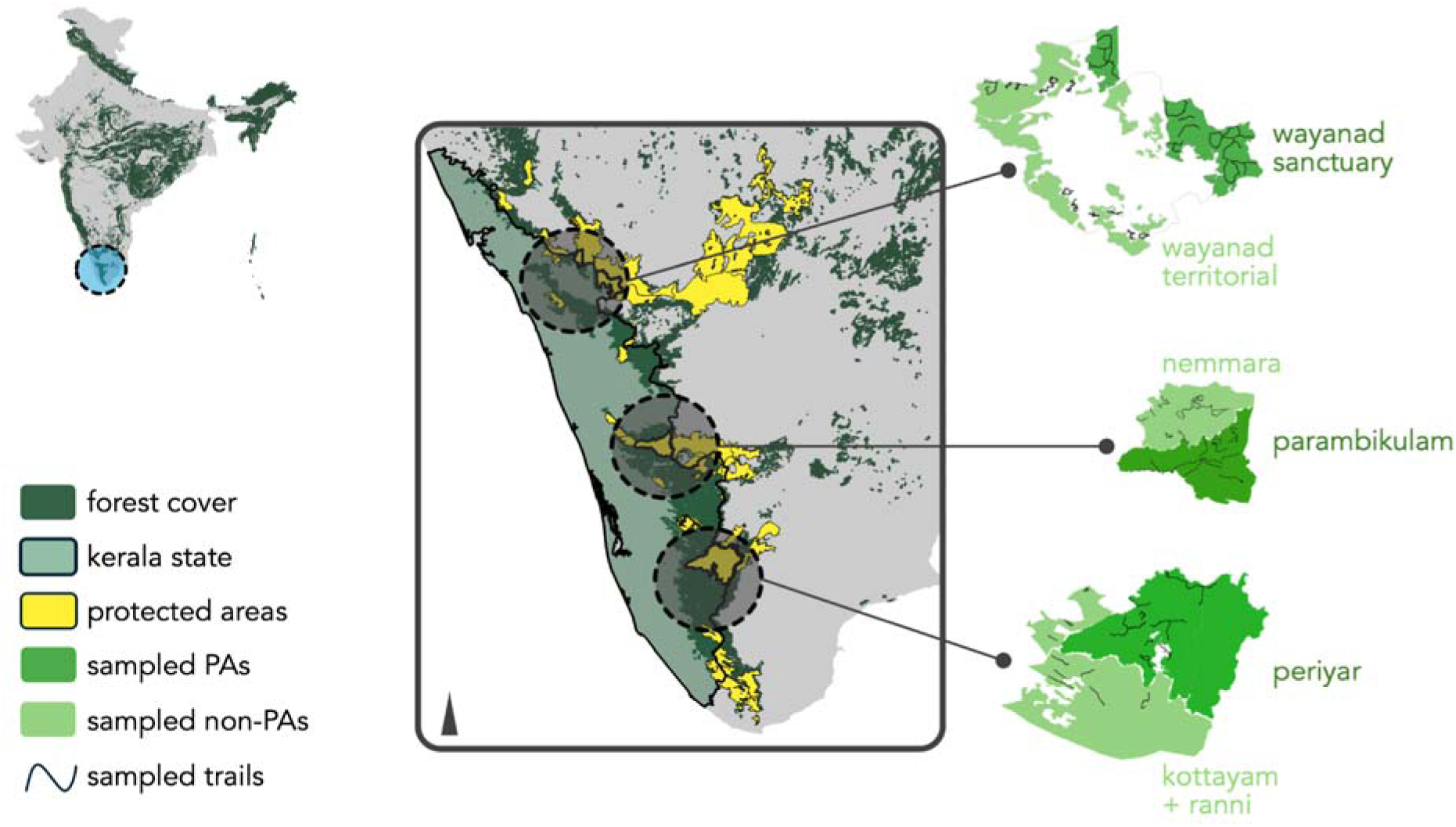
Study locations for field surveys in the state of Kerala in India’s Western Ghats. The map shows three Protected Areas: Wayanad Sanctuary, Parambikulam, and Periyar, and the surrounding unprotected forests and production agroforests in Territorial Divisions: Wayanad Territorial, Nemmara and Kottayam+Ranni Divisions. Black lines within the sites depict sampled survey routes (forest/plantation trails). Top-left: Location of the study area in India.

We surveyed 69 pre-determined routes in dhole habitats (forests and agroforests). These transects (3–18 km) followed forest/plantation trails, strategically placed to maximize spatial coverage of the study sites (**Fig. 1**). Each transect was divided into 100m segments and surveyed 1–6 times. Dhole scats were identified based on size, shape, smell, deposition style, and location (latrine sites; Andheria et al. 2007). Surveyors collected DNA from scats; only fresh scats (deposited within 1–2 days in direct sunlight or 2–3 days under canopy shade) were sampled. Two separate swabs were drawn on each piece of scat and stored in lysis buffer solution (Longmire et al. 1997; Ramón-Laca et al. 2015). For every sample, surveyors recorded geographic coordinates and descriptive details (condition, position, number of scats in the pile, and other remarks). Surveys were conducted in the post-monsoon months from October to May, typically over a three–five-week period in each site. In this paper, we present results from 2019 to 2023. In 2019 and 2022, surveys were conducted only in Wayanad Sanctuary. In 2023, the surveys were expanded to include Wayanad Sanctuary, Wayanad Territorial, Parambikulam, Nemmara, Periyar and Kottayam+Ranni.

### 2.4. Laboratory and analytical methods

We adapted the Single Nucleotide Polymorphism ‘SNP’ genotyping workflow standardized by Srivathsa et al. (2021) for identifying individual dholes from fecal DNA. Briefly, DNA is first extracted from the fecal samples using Qiagen DNA Extraction kit following the manufacturer’s protocol (with slight modifications). The next step involves a multiplex PCR where individual extracts are amplified using a panel of SNP-specific primers. The amplified products are then diluted and subjected to an indexing-PCR to attach unique barcodes to each product. Finally, these PCR products are pooled, purified and sequenced using an Illumina MiSeq kit. Srivathsa et al. (2021) developed primer pairs for 150 SNP loci with varying amplification success. Before processing our samples, we sought to retest the efficacy of this panel using a subset of our extracts (n = 32) and two dhole tissue samples. In this process, 54 primer pairs did not yield any amplification across the test samples; we therefore retained primer pairs targeting 96 SNPs for processing all samples in this study. The sequenced reads were analyzed further using standard bioinformatics pipelines: trimming the raw reads by removing barcode sequences using TrimGalore! (Krueger, F) with the following options: “quality 30”, “phred33”, “stringency 5”, “length 30” and “max_n 1”, mapping them onto a dog reference genome using BWA (Li 2013) and sorting using SAMtools (version 1.9; Li et al. 2009), filtering for quality control (mapping quality ≥30) and SNP calling using BCFtools. GATK (DePristo et al. 2011) was used to retain genotypes with a genotype quality ≥30 and depth ≥15; from this set of SNPs, VCFtools was used to retain only SNPs that had a minimum site quality of 30. This was followed by the estimation of pair-wise genetic relatedness among samples using PLINK (version 1.9; Purcell et al. 2007); the relatedness scores range from 0 (samples belong to unrelated individuals) to 1 (samples belong to the same individual; see **Supplementary File 2**). For a subset of the samples, we estimated relatedness between the replicate swab-draws (which can often be <1 when there are missing or poor-quality SNPs) to set a threshold (≥0.80) for assigning samples to individual dholes from the broader dataset. In other words, all sample-pairs from the entire sample pool whose relatedness score was ≥0.80 were assigned to the same individual (**Supplementary File 2**).

For each site, we first defined the state-space considering a buffer of 10km from the outermost sampled segments. Within this state-space, we overlaid a grid-array of 0.25 sq.km cells. Cells that contained ≥80% of dhole habitat (forest or agroforest) were retained as habitat ‘1’ cells; all others were removed and treated as non-habitat ‘0’ areas. The centroids of the retained cells collectively formed the mask layer, and a subset of these centroids (cells where the survey routes intersected) formed the detector array. Dhole detections were linked to the nearest detector to create individual-wise histories of spatial captures and recaptures. We used the classical likelihood-based SECR model (see Borchers and Efford 2008; Efford et al. 2009) to estimate baseline detection probability (g0), movement parameter associated with spatial scale of detection (σ) and population density (D). We first examined population densities within Wayanad Sanctuary across three years (2019, 2022, 2023) using the base SECR model with no covariates (g0∼1,σ∼1, D∼1). For 2023, we ran three analyses, one for each PA and the adjacent TD together, while modeling σ and D to vary based on location (PA versus TD; covariate model: g0∼1, σ ∼protected area, D∼protected area). For all the analyses, we compared the exponential (exp), half-normal (hnm) and hazard half-normal (hhn) models and chose the best one based on AICc scores. We note here that the hhn model estimates basal encounter rate (λ0) instead of probability (g0); in the text, tables and supplementary files, the ‘g0’ for hhn models need to be interpreted as the basal encounter rate (λ0). SECR analyses were implemented using the ‘secr’ package in R (R Development Core Team 2026).

We calculated abundance (N) differently for inside versus outside PAs. For inside the PAs, we multiplied the estimated density with the extent of the PA administrative boundary. Outside PAs, the TDs were expansive with no clear administrative limits. Here, we applied a buffer of √5.99 ✕ σ to the outer bounds of the sampled routes (Royle et al. 2013; Srivathsa et al.2021) and treated these polygons as ‘Effective Sample Areas’ or ESAs. For each TD, we multiplied the estimated density with the extent of the corresponding ESA. In sum, abundance in PA = D_pa_ ✕ administrative area of PA; abundance in TD = D_npa_ ✕ ESA. Since we had multi-year data from Wayanad Sanctuary for three time periods (2019, 2022, 2023), we undertook a cursory examination of localized movements of individuals, and an exploratory assessment of apparent survival by comparing inter-annual recaptures against the estimated population sizes from SECR; the data were too sparse to implement formal open models such as Cormack-Jolly-Seber, Open SCR, or its variants (Lebreton et al. 1992; Gardner et al. 2018).

## 3. Results

In terms of the Forest Department personnel conducting field surveys and co-collecting samples with the research teams, we had an average of 46 staff members participating per site–year from 2019 to 2025. Here, our calculations include numbers for Wayanad Sanctuary with Wayanad Territorial combined under a common bracket ‘Wayanad’, Parambikulam with Nemmara ‘Parambikulam’, and Periyar with Kottayam+Ranni ‘Periyar’. Overall, the average number of personnel involved per year was the lowest in Wayanad (∼28 staff members), followed by Parambikulam (∼47) and Periyar (∼73). Across sites–years, the number of Watchers (Forest Watchers, Tribal Watchers, EDC members, NMR Watchers/Elephant Watchers) was much higher than BFO/SFO (90% versus 10%; **Fig. 2**).

**Figure 2.**
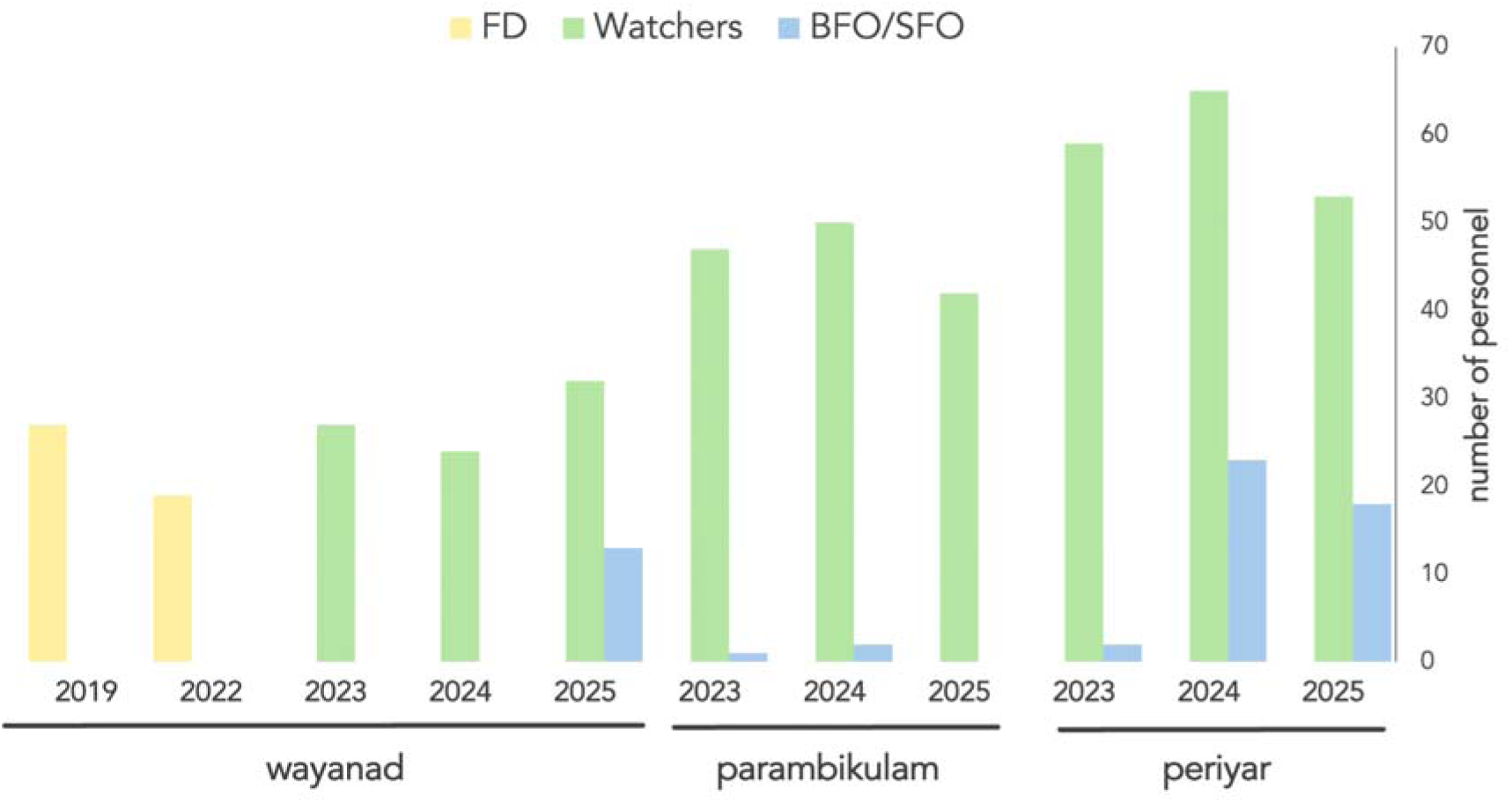
Number of Forest Department personnel involved in conducting field surveys and co-collecting fecal DNA samples (2019–2025). The personnel include Watchers (Forest Watchers, Tribal Watchers, Eco Development Committee ‘EDC’ members, Nominal Muster Roll ‘NMR’ Watchers/Elephant Watchers), Beat Forest Officers (BFO) and Section Forest Officers (SFO). Data are combined for Wayanad Sanctuary–Wayanad Territorial as ‘Wayanad’, Parambikulam–Nemmara as ‘Parambikulam’, and Periyar–Kottayam+Ranni as ‘Periyar’. In 2019 and 2022, ‘FD’ represents the combined number of Watchers and BFO/SFO for Wayanad where we did not make the distinction in designations while recording personnel particulars.

From 2019 to 2025, we received an average of ∼90 internship applications per year; of these we typically selected three–five interns per site–year. We were able to obtain responses from 29 of 30 former interns (RR: 97%) for our feedback survey. By design, every cohort was selected ensuring gender-balance in the entire research team; the respondents therefore consisted of 63% female interns. For most (41%), this project was their first field-based research experience; for 38% of them, this was their second or third such stint. In the time that elapsed since their internship, over half of them (52%) ended up as working professionals in wildlife-related fields, or are currently pursuing higher studies in wildlife-related fields (31%). The average rating based on a 5-point Likert scale was 4.6 (1.04SD) for four of the five questions regarding (i) increase in familiarity with field research, (ii) the advantages of gender-balanced teams, (iii) the training provided prior to the surveys and (iv) the extent to which their participation in this project increased their interest toward pursuing a career in wildlife-related research (see **Supplementary File 1**). Only one question consistently got a lower rating: the stipend offered did not appear to be a key motivation for their decision to join the project (average score = 2.35, 1.29SD; **Fig. 3**).

**Figure 3.**
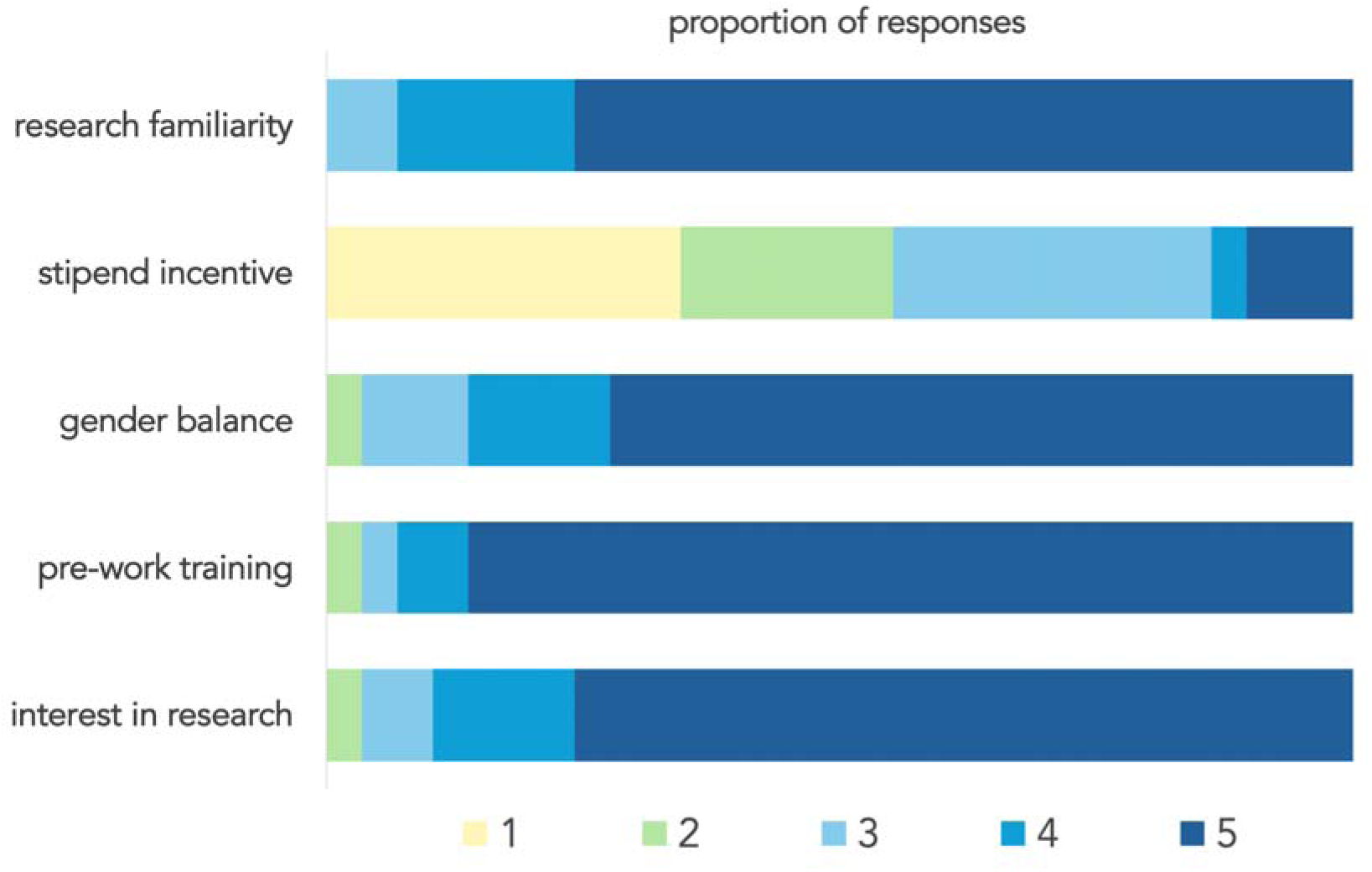
Responses elicited from citizen-volunteers (n=29) as part of the project feedback form. Former interns were asked to rate the extent to which they agreed with statements regarding five attributes: *the project increased their understanding of scientific research* (research familiarity), *the stipend offered influenced their decision to participate* (stipend incentive), *it was beneficial to have gender-balanced teams* (gender balance), *the training provided prior to the surveys was useful* (pre-work training), and *the project increased their interest in pursuing wildlife-related research/career* (interest in research). Likert scale 1–5: 1=Strongly disagree, 2=Disagree, 3=Neutral, 4=Agree, 5=Strongly agree. The full survey form with associated details is provided in Supplementary File 1.

A summary of the survey walk effort in each site–year, number of fecal samples collected, genotyping success rate, and the number of unique dhole individuals identified are presented in **Table 1**. In Wayanad Sanctuary (2019, 2022 and 2023), the estimated density (D) marginally increased from 15.67 (SE4.32) per 100 sq.km in 2019 to 19.01 (4.73SE) per 100 sq.km in 2023; but the respective 95% Confidence Intervals showed substantial overlap, indicating that the densities, and the population sizes (N = 57, 59 and 69), within the PA did not change significantly over the time period (**Fig. 4, top panel**; **Supplementary File 3**). Comparing across six sites in 2023, based on analyses performed for each PA and its adjacent TD together, the lowest estimated density was in Parambikulam (D per 100 sq.km = 9.73, 4.57SE), and highest in the adjacent TD of Nemmara (D per 100 sq.km = 29.23, 12.34SE). In Wayanad, the densities were somewhat similar in the Sanctuary versus the surrounding TD, in Parambikulam the density was higher outside the PA as compared to inside, and in Periyar, the density inside the PA was higher than outside (**Fig. 4, bottom panel**). Abundances ranged from 34 individuals inside Parambikulam to 196 individuals in Periyar (there would be some overlap of individuals within and outside PAs, because the ESAs intersect PA boundaries in some places). Model estimates of g0, σ and D, and the computed abundances (N) for all sites–years are in **Table 2**. The summed conditional activity-centre probability surface for detected individuals within the state-space, derived from the fitted SECR models, are mapped in **Figure 4**. Model comparisons and a compilation of estimates from all model outputs are provided in **Supplementary File 3**.

**Figure 4.**
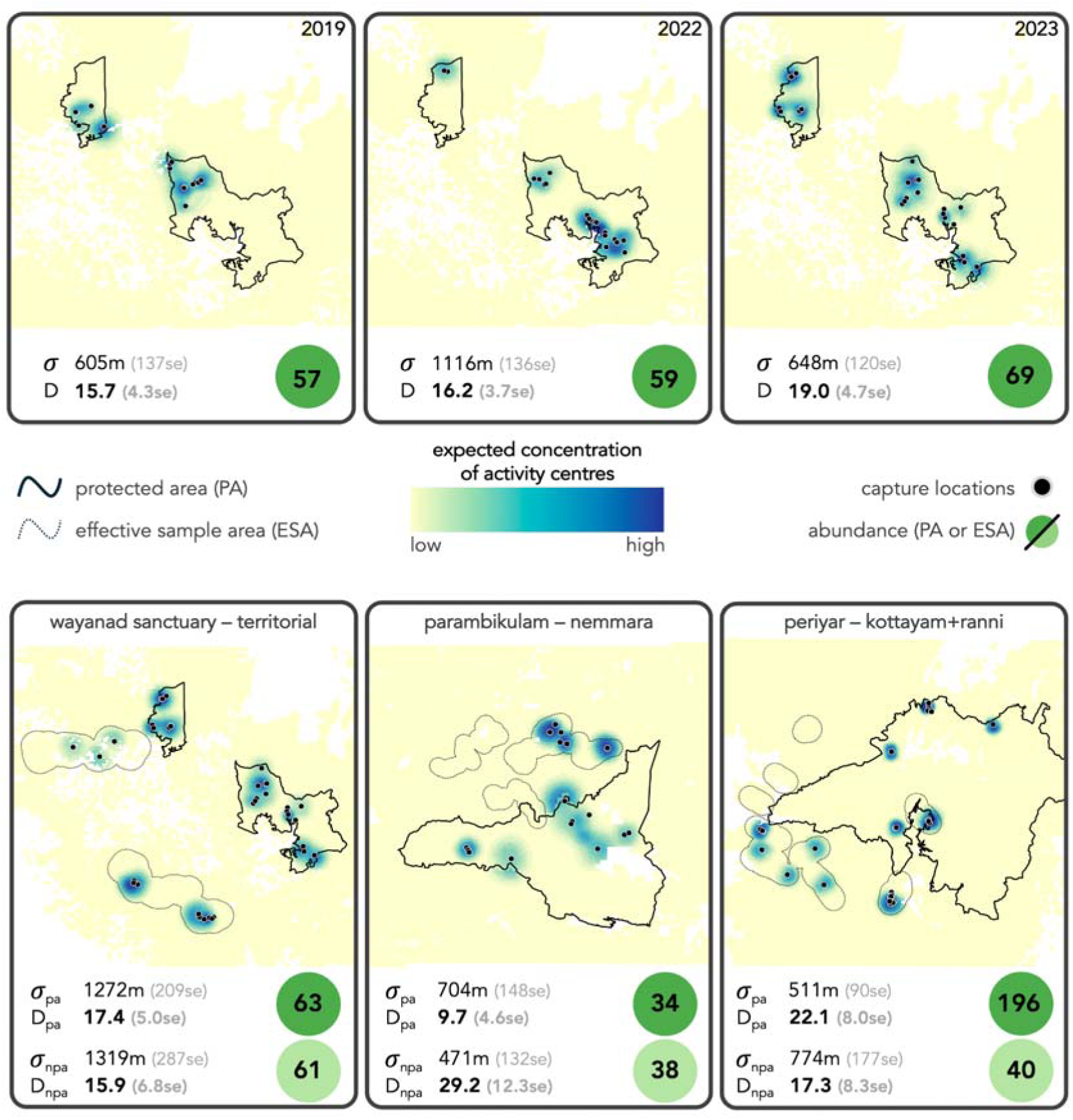
Summed conditional activity-centre probability surface for detected individuals within the state-space, derived from the fitted SECR models. Darker colors indicate higher expected concentration of activity centres. Top panel: Results from Wayanad Sanctuary over three time-steps (2019, 2022 and 2023); estimated sigma () and density (D) values are provided with corresponding standard errors in parentheses. Bottom panel: Results from 2023 in Wayanad Sanctuary–Wayanad Territorial, Parambikulam–Nemmara and Periyar–Kottayam+Ranni; sigma () and density (D) values estimated separately for inside Protected Areas (subscript ‘pa’) and Territorial Divisions outside Protected Areas (subscript ‘npa’); the numbers within circles at th bottom-right of each map are the estimated abundances, calculated as density ✕ PA size (for PAs) and density ✕ Effective Sample Area (for ESAs in TDs).

**Table 1.**
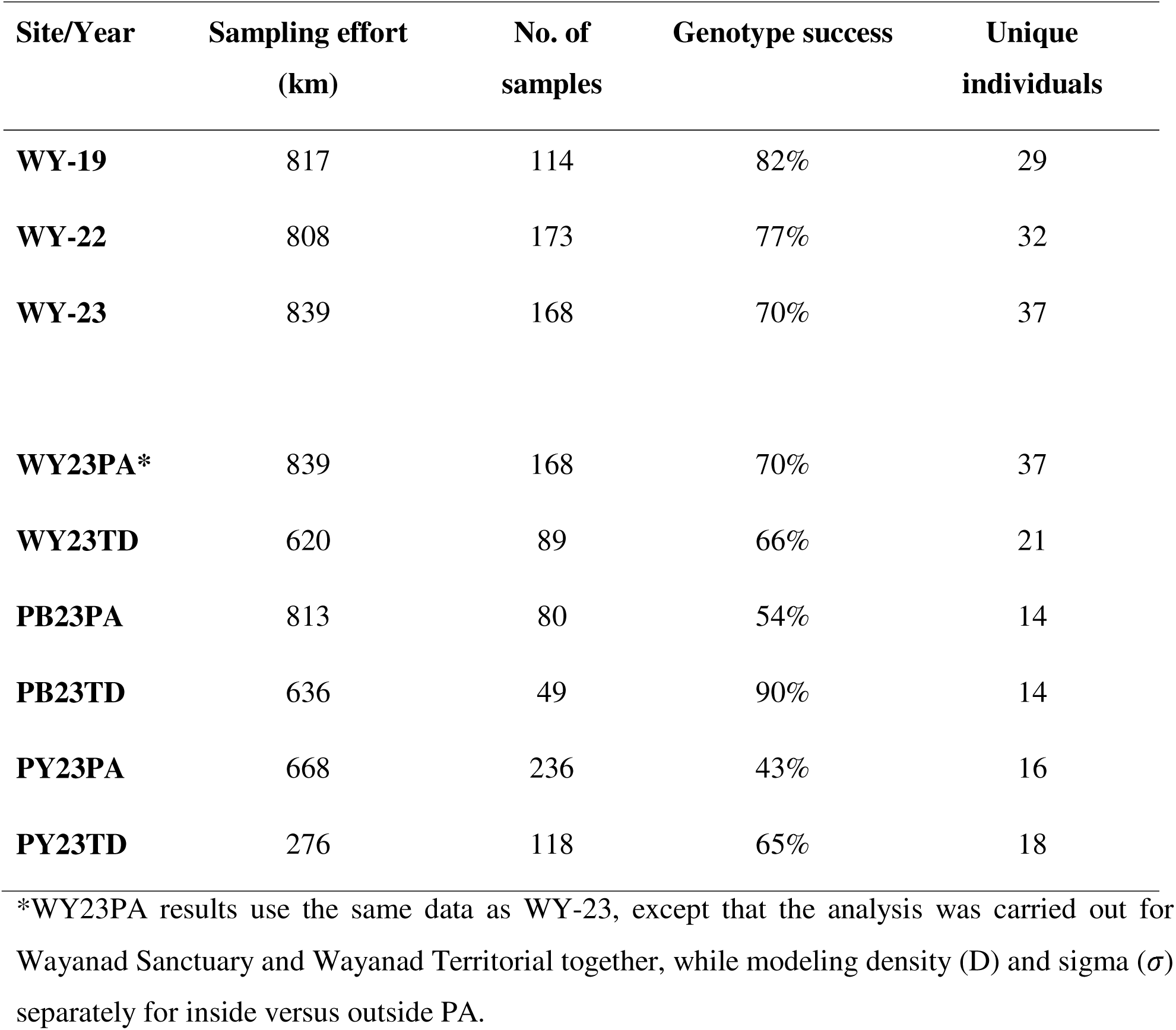
Summary of walk survey effort (km), number of field samples collected, genotype success (%) and the number of unique dhole individuals identified across all sites and years. The site–year codes refer to 3 years of Wayanad Sanctuary (WY-19: 2019, WY-22: 2022, WY-23: 2023), and 6 sites in 2023: Wayanad Sanctuary (WY23PA), Wayanad Territorial (WY23TD), Parambikulam (PY23TD). (PB23PA), Nemmara (PB23TD), Periyar (PY23PA) and Kottayam+Ranni.

| Site/Year | Sampling effort<br>(km) | No. of<br>samples | Genotype success | Unique<br>individuals |
| --- | --- | --- | --- | --- |
| **WY-19** | 817 | 114 | 82% | 29 |
| **WY-22** | 808 | 173 | 77% | 32 |
| **WY-23** | 839 | 168 | 70% | 37 |
| **WY23PA\*** | 839 | 168 | 70% | 37 |
| **WY23TD** | 620 | 89 | 66% | 21 |
| **PB23PA** | 813 | 80 | 54% | 14 |
| **PB23TD** | 636 | 49 | 90% | 14 |
| **PY23PA** | 668 | 236 | 43% | 16 |
| **PY23TD** | 276 | 118 | 65% | 18 |

\*WY23PA results use the same data as WY-23, except that the analysis was carried out for Wayanad Sanctuary and Wayanad Territorial together, while modeling density (D) and sigma ( $\sigma$ ) separately for inside versus outside PA.

**Table 2.** Results for estimated basal encounter rate (g0), movement parameter (σ) and density (D) from SECR models for three years in Wayanad Sanctuary (2019, 2022, 2023) and six sites in 2023: Wayanad Sanctuary (WY23PA), Wayanad Territorial (WY23TD), Parambikulam (PB23PA), Nemmara (PB23TD), Periyar (PY23PA) and Kottayam+Ranni (PY23TD).

| | Model | $g_0$<br>(SE) | $\sigma$<br>(SE) | PA size<br>(km<sup>2</sup>) | ESA<br>(km<sup>2</sup>) | D (SE)<br>/100km<sup>2</sup> | N<br>(range) |
| --- | --- | --- | --- | --- | --- | --- | --- |
| **WY19** | exp | 0.12 (0.06) | 605 (137) | 362 | – | 15.67 (4.32) | 57 (41–72) |
| **WY22** | hhn | 0.03 (0.01) | 1116 (136) | 362 | – | 16.21 (3.72) | 59 (45–72) |
| **WY23** | exp | 0.09 (0.03) | 648 (120) | 362 | – | 19.01 (4.73) | 69 (52–86) |
| **WY23PA\*** | hnm | 0.02 (0.01) | 1272 (209) | 362 | – | 17.35 (4.95) | 63 (45–81) |
| **WY23TD** | hnm | 0.02 (0.01) | 1319 (287) | – | 385 | 15.86 (6.79) | 61 (35–87) |
| **PB23PA** | exp | 0.06 (0.02) | 704 (148) | 352 | – | 9.73 (4.57) | 34 (18–50) |
| **PB23TD** | exp | 0.06 (0.02) | 471 (132) | – | 131 | 29.23 (12.34) | 38 (22–54) |
| **PY23PA** | hhn | 0.10 (0.02) | 511 (90) | 885 | – | 22.13 (7.98) | 196 (125–266) |
| **PY23TD** | hhn | 0.10 (0.02) | 774 (177) | – | 232 | 17.29 (8.29) | 40 (21–59) |

\*WY23PA results use the same data as WY-23, except that the analysis was carried out for Wayanad Sanctuary and Wayanad Territorial together, while modeling density (D) and sigma ( $\sigma$ ) separately for inside versus outside PA (covariate model: $D \sim \text{protected area}$ , $\sigma \sim \text{protected area}$ ). Models: exp— exponential, hhn— hazard half-normal, hnm— half normal; Abundance (N) calculated as density × PA size (for inside PAs) and density × Effective Sample Area (for ESAs in TDs); the min–max range for N is calculated using $D-(SE)$ and $D+(SE)$ and multiplying these with PA size and ESA.

Examining individual captures across years in Wayanad Sanctuary (2019, 2022, 2023), we found one individual from 2019 recaptured in 2022; similarly, six individuals from 2022 were recaptured in 2023. There were no identified individuals that were common between 2019 and 2023. The extent of localized across-year movement of recaptured individuals averaged at ∼6.5km from 2019 to 2023 (range: 0.5–12km; **Supplementary File 4**); the average was 5.5km when only the individuals from 2022–23 were used, i.e., inter-annual movement in one year. As a cursory assessment of apparent survival, we compared inter-annual recaptures with the estimated population sizes from SECR. The proportion of individuals detected within each sampling year was approximately 51% in 2019 (29/57), 54% in 2022 (32/59), and 54% in 2023 (37/69). Using these as crude measures of detection, the single individual detected in both 2019 and 2022 gave an estimated three-year apparent survival of approximately 0.06 after adjusting for the 2022 detection fraction; expressed as an annual rate, this is 0.40. Similarly, the six individuals detected in both 2022 and 2023 correspond to an estimated annual apparent survival of 0.35 after adjusting for the 2023 detection fraction. The two independent intervals gave somewhat similar annual apparent survival estimates (0.35–0.40). This consistency is suggestive, but the values should be regarded as exploratory; using the proportion of individuals detected relative to SECR-estimated abundance as a correction for partial detection is not a robust method for estimating apparent survival.

## 4. Discussion

Dhole individuals are morphologically indistinguishable from each other, the species lives in groups, and dholes perceivably occur at low local densities across much of their range. As a consequence, they have only been incidentally included in most multi-species or multi-year monitoring studies across South and Southeast Asia; in cases where the species was included, researchers have, at best, estimated occupancy probabilities (e.g., Phumanee et al. 2020; Chaudhary et al. 2022). Our work therefore holds significance on two fronts––we demonstrate how: (1) SNP-based individual identification combined with SECR models can offer a scaffolding to design dhole population monitoring programs across space and time in various parts of their geographic range; and (2) such monitoring can be sustained when implemented as a multi-agency or cross-institutional undertaking while also building local capacity.

### 4.1. Population parameters and local ecological contexts

Making a reasonable assumption that Protected Areas have higher concentration of resources and lower anthropogenic pressures, we expected that g0 would be higher and σ would be lower inside PAs (Efford et al. 2016); consequently, density (D) would be higher within PAs compared to the landscape mosaic outside (Rather et al. 2021; Sharma et al. 2021). Our results did not fully align with this expectation; in fact, our overall results hint at the possibility of a panmictic metapopulation in this landscape, rather than a classical source–pseudo-sink system. Importantly, the high density in Nemmara appeared rather peculiar. For context, this site is an extension of the Valparai agroforest landscape in the neighboring state of Tamil Nadu; dholes occur at high densities in Valparai, with 5–6 resident packs and density of >20 per 100 sq.km (Pious et al. 2025; D. Ganguly et al., unpublished results). Viewed in tandem, we propose a few possible explanations: first, the high densities in Territorial Divisions are because of adequate prey availability *in conjunction* with low intraguild competition with tigers and leopards (Karanth and Sunquist 2000; Srivathsa et al. 2023; A. Srivathsa, unpublished data). It is likely that fewer but larger packs are exploiting this niche––which may be unique to the forest–agroforest mosaics of the Western Ghats, but does not hold true for unprotected forests outside PAs elsewhere in the country (Srivathsa et al. 2019, 2020a). Second, low conflict with humans in the landscape makes it conducive for dhole packs to thrive in shared agroforest habitats (Srivathsa et al. 2020c; Saravanan et al. 2026). Third, there may be a biogeographic barrier-induced accumulation of dhole populations to the north of Parambikulam (Nemmara) because of the Palghat Gap (see Biswas and Karanth 2021)––where conditions of sufficient prey, low competition and benign interactions with people coalesce to create an optimal zone of high densities.

The limited number of individuals and the relatively short duration of the study (four years, 2019–2023) notwithstanding, our results showed some evidence for localized movement in Wayanad Sanctuary. It remains unclear if entire dhole packs shift territories, or if the observed movements arose from individuals emigrating and dispersing between packs (see Venkataraman 1998). Application of Open SCR models (e.g., Gardner et al. 2018) can help better delineate patterns of local movement across years. In addition, detailed examination of pair-wise relatedness of individuals can enable a more conclusive assessment (Meiring et al. 2022; vonHoldt et al. 2024); unfortunately, our SNP numbers and amplification depth did not allow for making assessments of relatedness scores to tease these aspects apart. Apparent survival offers a glimpse into the underlying biological process that determines population growth rate. In solitary large carnivores, growth rate is sustained by high rates of adult survival (>60–70%; see Karanth et al. 2006; Steinmetz et al. 2025). Rudimentary calculations from Wayanad Sanctuary––where dhole populations remained stable over four years––place apparent survival at 35–40%; this corroborates demographic rates documented in social carnivores, where fecundity and pup/juvenile survival (conditioned on survival of the breeding adults), rather than overall adult survival, plays a more substantive role in shaping population growth rates (Watts and Holekamp 2009; Davies-Mostert et al. 2015).

### 4.2. Practical and analytical considerations

The advent of SNP genotyping has revolutionized the application of genomics in population ecology (Helyar et al. 2011). Although the initial investment can be substantial, most are one-time costs, and yield high return on investment (Natesh et al. 2019). Srivathsa et al. (2021) reported a one-time cost of up to 25USD per primer pair and a per-sample sequencing cost of 8USD (1825INR and 580INR, respectively). However, primers are sensitive to repeated freeze–thaw cycles, which may compromise nucleotide integrity and reduce genotyping success (Davis et al. 2000). This is relevant for long-term studies such as ours, where primer solutions stored at −20 or −80°C need to be repeatedly thawed during aliquoting or get inadvertently thawed during freezer maintenance or malfunction. In such cases, additional investments may be required to replace degraded primers and maintain genotyping success. We note that Srivathsa et al. (2021) used a larger panel of 150 SNPs and achieved 83% genotyping success. We used a subset of these SNPs and observed variable success (43–90%) across sites and years. Srivathsa et al. (2021) also used the multiple observation process ‘MOP’ SCR approach that integrates detections of marked individuals and unmarked detections (indirect signs) to fully utilize all available field data (Tourani et al. 2020). Although the MOP–SCR model offers greater analytical sophistication, we relied instead on the classical likelihood-based SECR (Borchers and Efford 2008), whose strength lies in its relative simplicity and ease of implementation. Despite these differences, density estimates for Wayanad Sanctuary in 2019 between the two studies remained consistent.

In terms of practical considerations for field survey design, group-living species may show varying levels of heterogeneity in detection owing to variations in social dynamics, i.e., movement patterns, group size, cohesion, and aggregation, potentially rendering standard SECR models unreliable (Cubaynes et al. 2010; Emmet et al. 2021). Since dholes occur in moderately cohesive groups (average pack size ranging from 5–15 individuals; Srivathsa et al. 2017, 2020c; Saravanan et al. 2026), this concern is somewhat ameliorated (see López-Bao et al. 2018; Bischof et al. 2020). Within the SECR approach, population estimation using genetic individual identification requires repeated sampling for animal feces along fixed trails. Hence, detections are seldom fixed in space and can occur in variable locations along these trails. This can then be fitted into the search-encounter framework to model activity centres and density (Royle et al. 2011). An alternative approach is to overlay a grid network over the sampling area and snapping detections to the nearest grid-cell centroid (where these centroids act as pseudo-detectors); this is especially useful when sampling does not follow a fixed design (Russell et al. 2012). We leveraged this *post hoc* grid-fitting approach because (1) it optimizes for processing time and is easier to implement for wildlife managers, (2) our grid-spacing of 500m is less than 1.5 ✕ σ, reducing chances of estimation bias (Milleret et al. 2018), and (3) it allowed us to align this analysis of dhole data with other ongoing assessments focused on their co-predators, prey species and human impacts in the same landscape.

### 4.3. Citizen-volunteers as assets in population monitoring

Citizen-volunteers can help scale-up the scope of research projects while rationalizing personnel involvement. Many monitoring programs have thus been able to accrue massive volumes of data, enabling researchers to undertake sophisticated ecological assessments (Chandler et al. 2017; Freeman et al. 2022). In return, citizen-volunteers stand to benefit from practical exposure to wildlife research, fieldwork, and conservation practice (Newman et al. 2003; Forrester et al. 2017; Phukan et al. 2026). In our study, many of the interns were gaining their first or early field-based research experience. Our feedback survey provided some insight into the broader value of involving citizen-volunteers: many interns had limited prior experience, but several of them subsequently moved into wildlife-or conservation-related work or higher studies; their responses indicated that the training and research exposure gained were important aspects of their participation. Given that most PAs in India are highly restrictive in terms of citizen-access (besides tourism; Velho et al. 2012), we believe our monitoring framework helped foster *democratizing* field science in a small yet meaningful manner. Citizen-volunteer participation in conservation research also raises concerns about how it is valued and recognized––unpaid work can become exploitative and limit *who* is able to access these opportunities (Fournier and Bond 2015). In our work, the stipend offered did not seem to be a major motivation. However, this does not negate the importance of compensation; it remains relevant for acknowledging labor contributions and reducing financial barriers to participation. Further, having volunteers with different experiences, skills and perspectives can help complement each other during field-based monitoring (Aristeidou et al. 2021). Preferentially recruiting interns from the study landscape was extremely useful for us, as they were instrumental in coordinating field logistics. Balancing this with interns recruited from different regions, and maintaining gender balance, brought together people with diverse life experiences and skill sets. Such collaborative approaches could be useful for projects in the tropics, where sustained field effort and trained personnel can be invaluable assets for locally-empowered long-term biodiversity monitoring.

### 4.4. Iterative knowledge-sharing with multiple stakeholders

State-run institutions can play a key role in monitoring by facilitating permits and field logistics, and contributing knowledge (O’Connor et al. 2021). For research to inform management, relationships between researchers and managers should be established early by building trust and synergizing priorities (Merkle et al. 2019). Our repeated annual surveys fostered sustained engagement with department personnel, creating familiarity, continuity, and reciprocal understanding. The value of such work depends partly on whether knowledge and capacity can be transferred to existing institutions. We therefore designed field protocols, data entry–validation–archival procedures, laboratory workflows, and analytical approaches to facilitate seamless knowledge transfer to managers after project completion. To enable information flow, periodic reports were shared across departmental levels, beyond permit requirements. Regular training and interactions with ground staff strengthened this exchange; similar structured, tailored knowledge-transfer sessions will be implemented till the end of the current project timeline (2027), with opportunity for keeping open communication channels and expert inputs beyond the stipulated project end-date. Such iterative communication can bridge gaps between scientific and practitioner knowledge (Hulme 2014). Since frontline monitoring in India’s forests focuses primarily on tigers and elephants, these sessions provided avenues to introduce dhole ecology to managers and could support integration of dhole monitoring into existing systems such as M-STrIPES (Jhala et al. 2013). Next, dhole occurrence in agroforests adjoining PAs also necessitated building relationships with plantation managers and private landowners for landscape-scale monitoring. Alongside reports and meetings, posters and infographics in local languages, shared through WhatsApp, provided a low-cost conduit linking researchers, Forest Department personnel, and plantation managers. Finally, engagement with the IUCN Dhole Working Group enabled extension of this approach across the species’ range. Kao et al. (2020) identified collaborative networks, data compilation and sharing, and knowledge-gap identification as the Working Group’s key priorities. Sharing monitoring protocols and practical experience through this network is enabling locally tailored implementation, which can contribute toward future range-wide assessments, prioritization of dhole conservation landscapes, and national dhole conservation plans.

## 5. Conclusion

We acknowledge that Random-Encounter-Model-based estimates of population size using low-cost camera-trap data, like Space-To-Event models, can be of immense utility for dholes (Punjabi et al. 2022); these approaches are valuable for establishing snapshot population baselines across dhole range. However, individual ID-based approaches cannot be replaced––they are far more powerful and open pathways for applying tools to understand demographic parameters (e.g., Duangchantrasiri et al. 2016; Harihar et al. 2020) and tracking individuals over time (e.g., Kasper et al. 2026). Our results and monitoring framework hold promise, and we are cautiously optimistic on the prospects of estimating population vital rates, growth rates, and trends, especially the tentative contrasts in density variation inside versus outside PAs over time, that the data from 2024 onwards may reveal. Looking ahead, we envision the development of dynamic metapopulation models that incorporate population sizes, vital rates, anthropogenic stressors, prey species distribution, co-predator competition and connectivity considerations for dholes across the heterogenous Western Ghats mosaic landscape. Our work underscores how long-term studies are extremely important, but also entail significant logistical, financial, infrastructural, and human-power-related barriers (Lindenmeyer et al. 2012; Rafiq et al. 2024). From our own experience, this study was possible because of collaboration between institutions with equipment and expertise, the Forest Department with human power and local infrastructure, a large team of researchers, interns, and departmental ground staff, and a funding model that was part-supported by the Indian Government and in part through an international funding agency. Together, these partnerships and sustained investments will be critical for translating long-term research undertakings into robust evidence for conserving dhole populations across their range.

## Acknowledgements

We express gratitude to the Kerala State Forest Department for providing research permits (ref: WL10-28201/2018) and offering support in carrying out the study. In particular, we remain grateful to multiple PCCF/CWWs of Kerala: S. Kumar, B. Thomas, G. Singh and D. Jayaprasad for facilitating the project. The field work was possible because of proactive and relentless support from N. K. Ajayaghosh, S. Money, and all the field officers, guards, watchers and members of the Eco-Development Committee who conducted the field surveys. We thank S. Sharma, A. Sureshbabu, K. Thottathil, M. S. Arjun, S. Kotian, M. A. Agnishikhe, A. Das, T. Kothawalla, K. Chauhan, A. Simon, P. Anjana, A. Edayatil, A. Pious, A. Ahmed, V. Thavara, A. George, V. Pai, H. S. Thasmai, H. Sahal, E. A. Tom, R. Rohan, C. P. Harsha, J. Rozario, S. Ravi, M. A. Shaikh, E. B. Aswathy, P. Vishnu, J. Mathew, A. Raju, E. D. Sethulakshmi, J. Vedaanthagan, S. M. Mathew, K. Sreelakshmi, N. C. Akshaya, O. G. Adirsha, H. Deva, S. Nandakumar and A. Shibu for assistance with field surveys. We thank R. G. Rodrigues, A. Zachariah, R. W. Taylor, M. Natesh, A. Khan, H. Chhattani, B. V. A. Prasad, S. Darshan, V. Sagar, P. Praveen, I. Sinha, A. Chandramouli, V. Paynter and J. Manazhi for all their help in procuring samples, generating data, and conducting laboratory work and analyses. We acknowledge A. Pandit, C. P. Lakshminarayanan, S. Raj and K. Virbhadra from the Next Generation Genomics Sequencing, Centre for Cellular And Molecular Platforms (C-CAMP) facility for the Illumina sequencing runs, the NCBS Scientific Computing for high-performance cluster supported under project no. 12-R&D-TFR-5.04-0900 by the Department of Atomic Energy, Government of India, and P. Dey from the NCBS Collections Facility and NCBS Wildlife Program for providing working space. We are grateful to Wildlife Conservation Society–India, National Centre for Biological Sciences–TIFR and the Rufford Foundation for funding the 2019 study. In the subsequent years (2022–2025), the project received funding from the Conservation, Food and Health Foundation. A.S. was supported by the University of Florida (Gainesville, USA) and the Wildlife Conservation Network in 2019, and by the Department of Science and Technology–Government of India’s Innovation in Science Pursuit for Inspired Research (INSPIRE) Faculty Award from 2021 onwards.

## CRediT authorship contribution statement

**Arjun Srivathsa:** Writing – review & editing, Writing – original draft, Visualization, Validation, Software, Methodology, Investigation, Funding acquisition, Formal analysis, Conceptualization. **Mayank Shukla:** Writing – review & editing, Writing – original draft, Validation, Software, Methodology, Investigation, Formal analysis. **Pooja Saravanan:** Writing – review & editing, Writing – original draft, Validation, Investigation. **Divyajyoti Ganguly:** Writing – review & editing, Writing – original draft, Validation, Software, Methodology, Investigation, Formal analysis. **Midhun Mohan:** Writing – review & editing, Validation, Software, Methodology, Investigation, Formal analysis. **Uma Ramakrishnan:** Writing – review & editing, Validation, Methodology, Investigation, Funding acquisition, Supervision, Conceptualization.

## Declaration of competing interest

The authors declare that they have no known competing financial interests or personal relationships that could have appeared to influence the work reported in this paper.

## Data Availability Statement

The raw data used in the analyses will be deposited on a public repository upon manuscript acceptance.

## Supplementary Files

**Supplementary File 1.** Questionnaire survey form sent to citizen-volunteers (former interns) to elicit information regarding their experience participating in the project and its impacts on their subsequent professional trajectories.

**Supplementary File 2.** Summary of samples collected, the number of SNPs, missingness criteria, genotype success (%), unique individuals identified; distribution of pair-wise genetic relatedness among all sample pairs across Wayanad Sanctuary, Wayanad Territorial, Parambikulam, Nemmara, Periyar and Kottayam+Ranni from 2019 to 2023.

**Supplementary File 3.** Model comparisons, outputs and detailed list of parameters estimate from all the SECR model-fits for data on dhole encounters from 2019 to 2023, from Wayanad Sanctuary, Wayanad Territorial, Parambikulam, Nemmara, Periyar and Kottayam+Ranni.

**Supplementary File 4.** Dhole individuals recaptured across years (2019, 2022, 2023) in Wayanad Sanctuary and their localized movements between survey periods.

## Notes

### Competing Interest Statement

The authors have declared no competing interest.

