## Supplementary File 1 for "Leveraging collaborations between researchers, wildlife managers and citizen-volunteers to monitor endangered carnivore populations across space and time"

**Supplementary File 1. Questionnaire survey form sent to citizen-volunteers (former interns) to elicit information regarding their experience participating in the project and its impacts on their subsequent professional trajectories.**

We want to understand how participation in a long-term wildlife monitoring project contributes to the development of skills, knowledge, career opportunities, and professional growth among citizen-volunteers who were/are students and early-career researchers. This survey form is being sent out to all the people who interned with **[Project Name/Institution]** from 2019 to 2025.

Please note that your participation in this survey is very important to us. We assure you that all your responses will be anonymous and used only for research purposes.

ഒരു ദീർഘകാല വന്യജീവി നിരീക്ഷണ പദ്ധതിയിലെ പങ്കാളിത്തം, വിദ്യാർത്ഥികളും, പുതു ഗവേഷകരുമായ സന്നദ്ധപ്രവർത്തകരുടെ കഴിവുകൾ, അറിവ്, തൊഴിൽ അവസരങ്ങൾ, പ്രൊഫഷണൽ വളർച്ച എന്നിവയുടെ വികസനത്തിന് എങ്ങനെ സംഭാവന നൽകുന്നുവെന്ന് മനസ്സിലാക്കാൻ ഞങ്ങൾ ആഗ്രഹിക്കുന്നു. 2019 മുതൽ 2025 വരെ **[Project Name/Institution]** ഇന്റേൺ ചെയ്ത എല്ലാ ആളുകൾക്കും ഈ സർവേ ഫോം അയയ്ക്കുന്നുണ്ട്.

ഈ സർവേയിലെ നിങ്ങളുടെ പങ്കാളിത്തം ഞങ്ങൾക്ക് വളരെ പ്രധാനമാണെന്ന് ദയവായി മനസ്സിലാക്കുക. നിങ്ങളുടെ എല്ലാ പ്രതികരണങ്ങളും അജ്ഞാതമായിരിക്കുമെന്നും ഗവേഷണ ആവശ്യങ്ങൾക്കായി മാത്രം ഉപയോഗിക്കുമെന്നും ഞങ്ങൾ നിങ്ങൾക്ക് ഉറപ്പ് നൽകുന്നു.
______________________________________________________________________________

1. Before joining **[Project Name]**, how many wildlife-related projects/internships had you participated in?

**[Project Name]** മുമ്പ് നിങ്ങൾ എത്ര വന്യജീവി-അനുബന്ധ പ്രൊജെക്ടുകളുടെ/ഇന്റേൺഷിപ്പുകളുടെ ഭാഗമായിട്ടുണ്ട്?

☐ None (**[Project Name]** ആണ് ആദ്യത്തേത്)
 ☐ 1–2
 ☐ 3–5
 ☐ More than 5 അഞ്ചിലധികം

2. What best describes your current occupation or area of study?

   *Here, ‘wildlife-related’ includes ecology & evolution, wildlife, community-based conservation, environmental science, climate science, GIS and remote sensing, sustainability, forestry, natural resource management, environmental policy, restoration, naturalists/eco-tourism, environmental education, and related fields.*

നിങ്ങളുടെ ഇപ്പോഴത്തെ തൊഴിൽ /പഠനമേഖല എന്താണ്? ഏറ്റവും അനുയോജ്യമായത് തിരഞ്ഞെടുക്കുക.

*താഴെയുള്ളവയിൽ, "വന്യജീവി-അനുബന്ധം" എന്നത് പരിണാമം, സാമൂഹ്യാധിഷ്ഠിത പരിസ്ഥിതി സംരക്ഷണം, പരിസ്ഥിതി ശാസ്ത്രം, കാലാവസ്ഥ വിജ്ഞാനം, GIS & വിദൂര സംവേദനം, സുസ്ഥിരത, വനം, പ്രകൃതിവിഭവ മാനേജ്മെന്റ്, പരിസ്ഥിതി നയം, പുനഃസ്ഥാപനം, പ്രകൃതിശാസ്ത്രജ്ഞർ/പരിസ്ഥിതി ടൂറിസം, പരിസ്ഥിതി വിദ്യാഭ്യാസം, അനുബന്ധ മേഖലകൾ എന്നിവ ഉൾപ്പെടുന്നു.*

☐ Pursuing higher studies in a wildlife-related field (വന്യജീവി-അനുബന്ധ മേഖലയിൽ ഉപരിപഠനം നടത്തുന്നു)

 ☐ Working professional in a wildlife-related field (വന്യജീവി-അനുബന്ധ മേഖലയിൽ തൊഴിൽ ചെയ്യുന്നു)

☐ Pursuing higher studies in another unrelated field (മറ്റു മേഖലയിൽ ഉപരിപഠനം നടത്തുന്നു)

☐ Working professional in another unrelated field (മറ്റു മേഖലയിൽ തൊഴിൽ ചെയ്യുന്നു)

 ☐ Other (മറ്റുള്ളവ)

The **[Project Name]** has ensured that every intern gets paid a nominal stipend during the internship. The Project has also tried best to ensure that the teams are always gender balanced. Given this background, for the following questions, please indicate your level of agreement using the following scale:

1 = Strongly disagree
 2 = Disagree
 3 = Neutral
 4 = Agree
 5 = Strongly agree

എല്ലാ ഇന്റേൺകൾക്കും തുച്ഛമായ ഒരു തുക സ്റ്റൈപ്പന്റ് ആയി ലഭിക്കുന്നുവെന്നു **[Project Name]** ഉറപ്പു വരുത്തിയിട്ടുണ്ട്. പ്രോജക്ടിന്റെ പ്രവർത്തന സംഘങ്ങളിൽ ലിംഗാനുപാതതുല്യത പരിപാലിക്കാനും പരമാവധി ശ്രെമിച്ചിട്ടുണ്ട്. ഇവ

കണക്കിലെടുത്തുകൊണ്ട്, 3 മുതൽ 7 വരെയുള്ള ചോദ്യങ്ങൾ താഴെപറയുന്ന വിധം നിങ്ങളുടെ യോജിപ്പ് രേഖപ്പെടുത്തുക.

1 = ശക്തമായി വിയോജിക്കുന്നു

2 = വിയോജിക്കുന്നു

3 = നിഷ്പക്ഷം

4 = യോജിക്കുന്നു

5 = ശക്തമായി യോജിക്കുന്നു

3. “Participation in **[Project Name]** helped me familiarize myself with conducting scientific research, field work and lab/analytical techniques.”

   “**[Project Name]** പങ്കാളിത്തം എന്നെ ശാസ്ത്രഗവേഷണം, ഫീൽഡ് വർക്ക് എന്നിവയുടെ നടത്തിപ്പിനേപ്പറ്റിയും ലബോറോട്ടറി വിദ്യകൾ, അനാലിസിസ് എന്നിവയെപ്പറ്റിയും കൂടുതൽ പരിചയസമ്പന്നരാവാൻ സഹായിച്ചു.”

☐ 1 ☐ 2 ☐ 3 ☐ 4 ☐ 5

4. “The stipend offered by the project influenced my decision to take up this internship.”

“വാഗ്ദാനം ചെയ്ത സ്റ്റൈപ്പന്റ് ഇന്റേൺഷിപ്പിനു അവസരം സ്വീകരിക്കാനുള്ള എന്റെ തീരുമാനത്തെ സ്വാധീനിച്ചിരുന്നു.”

☐ 1 ☐ 2 ☐ 3 ☐ 4 ☐ 5

5. “Working in gender-balanced teams improved working conditions, functioning and learning experience.”

“തുല്യ ലിംഗാനുപാതമുള്ള ടീമുകളിൽ പ്രവർത്തിക്കുന്നത് ജോലി സാഹചര്യങ്ങളും, പ്രവർത്തനവും, പഠനാനുഭവവും മെച്ചപ്പെടുത്തി.”

☐ 1 ☐ 2 ☐ 3 ☐ 4 ☐ 5

6. “The training and orientation provided at the beginning of my internship adequately prepared me for my responsibilities.”

“ഇന്റേൺഷിപ്പിന്റെ തുടക്കത്തിൽ നൽകിയ പരിശീലനം എന്നെ എന്റെ ഉത്തരവാദിത്തങ്ങൾക്കായി വേണ്ടത്ര സജ്ജരാക്കി.”

☐ 1 ☐ 2 ☐ 3 ☐ 4 ☐ 5

7. "Participation in the project increased my interest in pursuing a higher studies/a career in a wildlife-related field.”

“ഈ പ്രോജെക്ടിൽ പങ്കാളിയായത് വന്യജീവി ഗവേഷണവും സംരക്ഷണവും എന്റെ തൊഴിലോ അല്ലെങ്കിൽ പാഠ്യവിഷയമായോ തിരഞ്ഞെടുക്കാനുള്ള താൽപ്പര്യം വർദ്ധിപ്പിച്ചു.”

☐ 1 ☐ 2 ☐ 3 ☐ 4 ☐ 5
