## Supplementary File 2 for "Leveraging collaborations between researchers, wildlife managers and citizen-volunteers to monitor endangered carnivore populations across space and time"

**Supplementary File 2. Summary of samples collected, the number of SNPs, missingness criteria, genotype success (%), unique individuals identified; distribution of pair-wise genetic relatedness among all sample pairs across Wayanad Sanctuary, Wayanad Territorial, Parambikulam, Nemmara, Periyar and Kottayam+Ranni from 2019 to 2023.**

**Table S1.** Summary of number of samples collected, overall SNPs, missingness criteria, number of samples passed, genotype success (%), and unique dhole individuals identified across all sites and years. The site-year codes correspond to Wayanad Sanctuary (WY-19: 2019, WY-22: 2022), Wayanad Sanctuary+Wayanad Territorial (WY-23: 2023), Parambikulam+Nemmara (PB-23: 2023), Periyar+Kottay	am+Ranni (PY-23: 2022).

| **Site–Year** | **No. of samples** | **Overall SNPs**  **(Out of 96)** | **Missingness criteria** | **No. of samples passed** | **Genotype Success (%)** | **Unique**  **IDs** |
| --- | --- | --- | --- | --- | --- | --- |
| WY-19 | 114 | 64 | 0.5 | 93 | 81.58 | 29 |
| WY-22 | 173 | 82 | 0.5 | 133 | 76.88 | 32 |
| WY-23 | 257 | 78 | 0.5 | 176 | 68.48 | 58 |
| PB-23 | 129 | 82 | 0.5 | 87 | 67.44 | 28 |
| PY-23 | 354 | 74 | 0.5 | 178 | 50.28 | 34 |

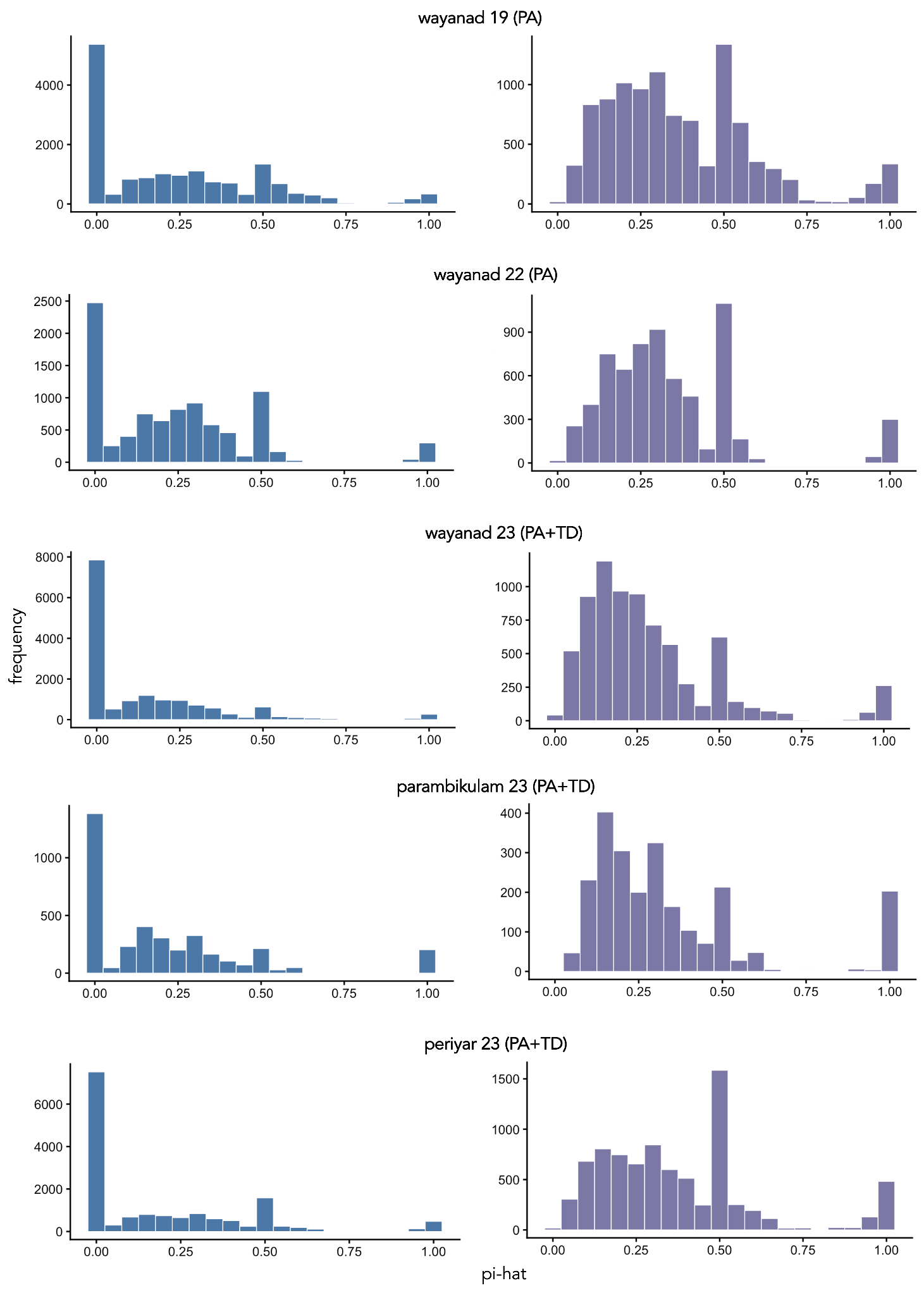
**Figure S1.** Distribution of pair-wise genetic relatedness (pi-hat) among all sample pairs across all sites and years (Wayanad Sanctuary 2019 and 2022, Wayanad Sanctuary and Wayanad Territorial 2023, Parambikulam and Nemmara 2023, Periyar and Kottayam+Ranni 2023). Left panel (blue): all relatedness scores including ‘zero’ (i.e., completely unrelated individuals). Right panel purple): relatedness scores excluding ‘zeroes’, plotted for easier visualization and interpretation of the non-zero scores (with prominent peaks at 0.25–half-sibs, 0.5–sibs, and 1–recaptures of same individual).
