## Supplementary File 3 for "Leveraging collaborations between researchers, wildlife managers and citizen-volunteers to monitor endangered carnivore populations across space and time"

**Supplementary File 3. Model comparisons, outputs and detailed list of parameter estimates from all the SECR model-fits for data on dhole encounters from 2019 to 2023, from Wayanad Sanctuary, Wayanad Territorial, Parambikulam, Nemmara, Periyar and Kottayam+Ranni.**

**Table S1.** Model comparisons based on AICc scores and weights for SECR models fitted to data from Wayanad Sanctuary (WY) in 2019, 2022 and 2023 (WY-19, WY-22, WY-23). The table includes comparisons between models with exponential (exp), half-normal (hnm) and hazard half-normal (hnm) detection functions.

|  | **Model** | **K** | **logLik** | **AICc** | **ΔAICc** | **AICc weight** |
| --- | --- | --- | --- | --- | --- | --- |
| **WY-19 (D~1, 𝜎~1, g0~1)** | | | | | | |
|  | exp | 3 | -178.714 | 364.387 | 0 | 0.7227 |
|  | hnm | 3 | -180.36 | 367.679 | 3.292 | 0.1394 |
|  | hhn | 3 | -180.37 | 367.7 | 3.313 | 0.1379 |
| **WY-22 (D~1, 𝜎~1, g0~1)** | | | | | | |
|  | hhn | 3 | -204.496 | 415.849 | 0 | 0.4567 |
|  | hnm | 3 | -204.503 | 415.863 | 0.014 | 0.4535 |
|  | exp | 3 | -206.123 | 419.103 | 3.254 | 0.0898 |
| **WY-23 (D~1, 𝜎~1, g0~1)** | | | | | | |
|  | exp | 3 | -212.913 | 432.553 | 0 | 0.447 |
|  | hnm | 3 | -213.391 | 433.508 | 0.955 | 0.2773 |
|  | hhn | 3 | -213.396 | 433.52 | 0.967 | 0.2757 |

K= number of parameters; logLik= log-likelihood

**Table S2.** Model comparisons based on AICc scores and weights for SECR models fitted to data from 2023 for Wayanad Sanctuary–Territoral (WY23 PA+TD), Parambikulam–Nemmara (PB23 PA+TD) and Periyar–Kottayam+Ranni (PY23 PA+TD). The table includes comparisons between models with exponential (exp), half-normal (hnm) and hazard half-normal (hnm) detection functions, for iteratively modified covariate models (parameter D or **𝜎** or both~protected area).

|  | **Model** | **K** | **logLik** | **AICc** | **ΔAICc** | **AICc weight** |
| --- | --- | --- | --- | --- | --- | --- |
| **WY23 PA+TD (D~1, 𝜎~1, g0~1)** | | | | | | |
|  | hnm | 3 | -328.0772 | 662.599 | 0 | 0.3738 |
|  | hhn | 3 | -328.0771 | 662.599 | 0 | 0.3738 |
|  | exp | 3 | -328.4701 | 663.385 | 0.786 | 0.2523 |
| **WY23 PA+TD (D~1, 𝜎~protected area, g0~1)** | | | | | | |
|  | hnm | 4 | -328.0772 | 664.909 | 0 | 0.3737 |
|  | hhn | 4 | -328.0771 | 664.909 | 0 | 0.3737 |
|  | exp | 4 | -328.4694 | 665.693 | 0.784 | 0.2525 |
| **WY23 PA+TD (D~protected area, 𝜎~1, g0~1)** | | | | | | |
|  | hnm | 4 | -328.0731 | 664.901 | 0 | 0.3721 |
|  | hhn | 4 | -328.073 | 664.901 | 0 | 0.3721 |
|  | exp | 4 | -328.4477 | 665.65 | 0.749 | 0.2559 |
| **WY23 PA+TD (D~protected area, 𝜎~protected area, g0~1)** | | | | | | |
|  | hnm | 5 | -328.063 | 667.28 | 0 | 0.3633 |
|  | hhn | 5 | -328.0632 | 667.28 | 0 | 0.3633 |
|  | exp | 5 | -328.3475 | 667.849 | 0.569 | 0.2734 |
| **PY23 PA+TD (D~1, 𝜎~1, g0~1)** | | | | | | |
|  | hhn | 3 | -205.879 | 418.557 | 0 | 0.4846 |
|  | hnm | 3 | -205.913 | 418.626 | 0.069 | 0.4682 |
|  | exp | 3 | -208.207 | 423.214 | 4.657 | 0.0472 |
| **PY23 PA+TD (D~1, 𝜎~protected area, g0~1)** | | | | | | |
|  | hhn | 4 | -203.647 | 416.673 | 0 | 0.4825 |
|  | hnm | 4 | -203.679 | 416.738 | 0.065 | 0.467 |
|  | exp | 4 | -205.903 | 421.186 | 4.513 | 0.0505 |
| **PY23 PA+TD (D~protected area, 𝜎~1, g0~1)** | | | | | | |
|  | hhn | 4 | -204.742 | 418.863 | 0 | 0.4845 |
|  | hnm | 4 | -204.777 | 418.933 | 0.07 | 0.4678 |
|  | exp | 4 | -207.061 | 423.501 | 4.638 | 0.0477 |
| **PY23 PA+TD (D~protected area, 𝜎~protected area, g0~1)** | | | | | | |
|  | hhn | 5 | -203.5705 | 419.284 | 0 | 0.4796 |
|  | hnm | 5 | -203.6012 | 419.345 | 0.061 | 0.4652 |
|  | exp | 5 | -205.7313 | 423.605 | 4.321 | 0.0553 |
| **PB23 PA+TD (D~1, 𝜎~1, g0~1)** | | | | | | |
|  | exp | 3 | -169.96 | 346.92 | 0 | 0.5781 |
|  | hnm | 3 | -170.966 | 348.933 | 2.013 | 0.2113 |
|  | hhn | 3 | -170.97 | 348.94 | 2.02 | 0.2106 |
| **PB23 PA+TD (D~1, 𝜎~protected area, g0~1)** | | | | | | |
|  | exp | 4 | -169.945 | 349.629 | 0 | 0.5623 |
|  | hnm | 4 | -170.887 | 351.513 | 1.884 | 0.2192 |
|  | hhn | 4 | -170.891 | 351.52 | 1.891 | 0.2185 |
| **PB23 PA+TD (D~protected area, 𝜎~1, g0~1)** | | | | | | |
|  | exp | 4 | -169.482 | 348.702 | 0 | 0.5671 |
|  | hnm | 4 | -170.443 | 350.625 | 1.923 | 0.2168 |
|  | hhn | 4 | -170.446 | 350.631 | 1.929 | 0.2161 |
| **PB23 PA+TD (D~protected area, 𝜎~protected area, g0~1)** | | | | | | |
|  | exp | 5 | -168.3333 | 349.394 | 0 | 0.4548 |
|  | hnm | 5 | -168.844 | 350.415 | 1.021 | 0.273 |
|  | hhn | 5 | -168.8463 | 350.42 | 1.026 | 0.2723 |

K= number of parameters; logLik= log-likelihood

**Table S3.** Key parameter estimates from fitted SECR models (g0, 𝜎, D), the associated Standard Errors (SE), 95% Confidence Intervals (CI); the table also includes calculated abundance in Protected Area (N_pa_), abundance in Effective Sample Area (N_esa_), and their respective min–max values (D–SE ✕ area, D+SE ✕ area). WY-19, WY-22, WY-23 are results for Wayanad Sanctuary in 2019, 2022, 2023.

|  | **Model** | **g0 (SE)** | **95% CI (g0)** | **𝜎 (SE)(m)** | **95% CI (𝜎)** | **PA size (km^2^)** | **D (SE) /100km^2^** | **95% CI (D)** | **N_PA_ (range)** | **ESA (km^2^)** | **N_ESA_ (range)** |
| --- | --- | --- | --- | --- | --- | --- | --- | --- | --- | --- | --- |
| **WY-19 (D~1, 𝜎~1, g0~1)** | | | | | | | | | | | |
|  | exp | 0.12 (0.06) | 0.04–0.28 | 605 (137) | 390–938 | 362 | 15.67 (4.32) | 9.22–26.63 | 57 (41–72) | 364 | 57 (41–73) |
|  | hnm | 0.04 (0.02) | 0.02–0.09 | 1097 (207) | 761–1581 | 362 | 14.68 (3.98) | 8.71–24.74 | 53 (39–68) | 537 | 79 (57–100) |
|  | hhn | 0.04 (0.02) | 0.02–0.09 | 1095 (207) | 759–1581 | 362 | 14.68 (3.98) | 8.71–24.74 | 53 (39–68) | 537 | 79 (57–100) |
| **WY-22 (D~1, 𝜎~1, g0~1)** | | | | | | | | | | | |
|  | hhn | 0.03 (0.01) | 0.02–0.05 | 1116 (136) | 880–1415 | 362 | 16.21 (3.72) | 10.41–25.26 | 59 (45–72) | 514 | 83 (64–102) |
|  | hnm | 0.03 (0.01) | 0.02–0.05 | 1119 (135) | 883–1417 | 362 | 16.22 (3.72) | 10.41–25.27 | 59 (45–72) | 515 | 83 (64–103) |
|  | exp | 0.07 (0.02) | 0.03–0.13 | 728 (110) | 542–977 | 362 | 16.19 (3.81) | 10.27–25.53 | 59 (45–72) | 374 | 61 (46–75) |
| **WY-23 (D~1, 𝜎~1, g0~1)** | | | | | | | | | | | |
|  | exp | 0.09 (0.03) | 0.04–0.16 | 648 (120) | 452–928 | 362 | 19.01 (4.73) | 11.76–30.72 | 69 (52–86) | 343 | 65 (49–81) |
|  | hnm | 0.03 (0.01) | 0.02–0.06 | 1061 (160) | 790–1424 | 362 | 18.55 (4.40) | 11.72–29.35 | 67 (51–83) | 494 | 92 (70–114) |
|  | hhn | 0.03 (0.01) | 0.02–0.06 | 1059 (160) | 788–1422 | 362 | 18.54 (4.40) | 11.72–29.34 | 67 (51–83) | 494 | 92 (70–114) |

**Table S4.** Key parameter estimates from fitted SECR models (g0, 𝜎, D), the associated Standard Errors (SE), 95% Confidence Intervals (CI); the table also includes calculated abundance in Protected Area (N_pa_), abundance in Effective Sample Area (N_esa_), and their respective min–max values (D–SE ✕ area, D+SE ✕ area) for SECR models fitted to data from 2023. WY23PA: Wayanad Sanctuary, WY23TD: Wayanad Territorial, PB23PA: Parambikulam, PB23TD: Nemmara, PY23PA: Periyar, PY23TD: Kottayam+Ranni.

|  | **Model** | **g0 (SE)** | **95% CI (g0)** | **𝜎 (SE)(m)** | **95% CI (𝜎)** | **PA size (km^2^)** | **D (SE) /100km^2^** | **95% CI (D)** | **N_PA_ (range)** | **ESA (km^2^)** | **N_ESA_ (range)** |
| --- | --- | --- | --- | --- | --- | --- | --- | --- | --- | --- | --- |
| **WY23PA (D~protected area, 𝜎~protected area)** | | | | | | | | | | | |
|  | hnm | 0.02 (0.01) | 0.01–0.04 | 1272 (209) | 923–1752 | 362 | 17.35 (4.95) | 10.03–30.01 | 63 (45–81) | 593 | 103 (74–132) |
|  | hhn | 0.02 (0.01) | 0.01–0.04 | 1270 (209) | 922–1750 | 362 | 17.35 (4.95) | 10.03–30.01 | 63 (45–81) | 565 | 98 (70–126) |
|  | exp | 0.06 (0.02) | 0.03–0.10 | 793 (151) | 548–1148 | 362 | 18.27 (5.36) | 10.40–32.09 | 66 (47–86) | 399 | 73 (51–94) |
| **WY23TD (D~protected area, 𝜎~protected area)** | | | | | | | | | | | |
|  | hnm | 0.02 (0.01) | 0.01–0.04 | 1319 (287) | 865–2011 | - | 15.86 (6.79) | 7.10–35.44 | - | 385 | 61 (35–87) |
|  | hhn | 0.02 (0.01) | 0.01–0.04 | 1317 (287) | 864–2007 | - | 15.87 (6.79) | 7.10–35.46 | - | 385 | 61 (35–87) |
|  | exp | 0.06 (0.02) | 0.03–0.10 | 905 (240) | 543–1508 | - | 13.70 (6.66) | 5.55–33.78 | - | 265 | 36 (19–54) |
| **PY23PA (D~protected area, 𝜎~protected area)** | | | | | | | | | | | |
|  | hhn | 0.10 (0.02) | 0.06–0.16 | 511 (90) | 363–719 | 885 | 22.13 (7.98) | 11.14–43.93 | 196 (125–267) | 212 | 47 (30–64) |
|  | hnm | 0.09 (0.02) | 0.06–0.15 | 515 (90) | 367–724 | 885 | 22.12 (7.99) | 11.13–43.95 | 196 (125–267) | 213 | 47 (30–64) |
|  | exp | 0.19 (0.05) | 0.11–0.32 | 359 (77) | 237–545 | 885 | 21.39 (8.20) | 10.36–44.20 | 189 (117–262) | 149 | 32 (20–44) |
| **PY23TD (D~protected area, 𝜎~protected area)** | | | | | | | | | | | |
|  | hhn | 0.10 (0.02) | 0.06–0.16 | 774 (177) | 497–1204 | - | 17.29 (8.29) | 7.09–42.17 | - | 232 | 40 (21–59) |
|  | hnm | 0.09 (0.02) | 0.06–0.15 | 782 (178) | 503–1214 | - | 17.24 (8.27) | 7.06–42.05 | - | 235 | 40 (21–59) |
|  | exp | 0.19 (0.05) | 0.11–0.32 | 587 (152) | 356–968 | - | 14.60 (7.23) | 5.83–36.57 | - | 168 | 25 (12–37) |
| **PB23PA (D~protected area, 𝜎~protected area)** | | | | | | | | | | | |
|  | exp | 0.06 (0.02) | 0.02–0.13 | 704 (148) | 469–1058 | 352 | 9.73 (4.57) | 4.05–23.36 | 34 (18–50) | 232 | 23 (12–33) |
|  | hnm | 0.02 (0.01) | 0.01–0.04 | 1178 (203) | 843–1646 | 352 | 9.23 (4.10) | 4.02–21.20 | 32 (18–47) | 340 | 31 (17–45) |
|  | hhn | 0.02 (0.01) | 0.01–0.05 | 1177 (202) | 842–1645 | 352 | 9.23 (4.10) | 4.02–21.20 | 32 (18–47) | 339 | 31 (17–45) |
| **PB23TD (D~protected area, 𝜎~protected area)** | | | | | | | | | | | |
|  | exp | 0.06 (0.02) | 0.02–0.13 | 471 (132) | 275–807 | - | 29.23 (12.34) | 13.22–64.65 | - | 131 | 38 (22–54) |
|  | hnm | 0.02 (0.01) | 0.01–0.04 | 762 (182) | 480–1208 | - | 29.08 (11.66) | 13.65–61.97 | - | 204 | 59 (35–83) |
|  | hhn | 0.02 (0.01) | 0.01–0.05 | 761 (182) | 479–1207 | - | 29.08 (11.66) | 13.65–61.97 | - | 203 | 59 (35–83) |
