## Supplementary figures and images for "Leveraging collaborations between researchers, wildlife managers and citizen-volunteers to monitor endangered carnivore populations across space and time"

### Supplementary File 4

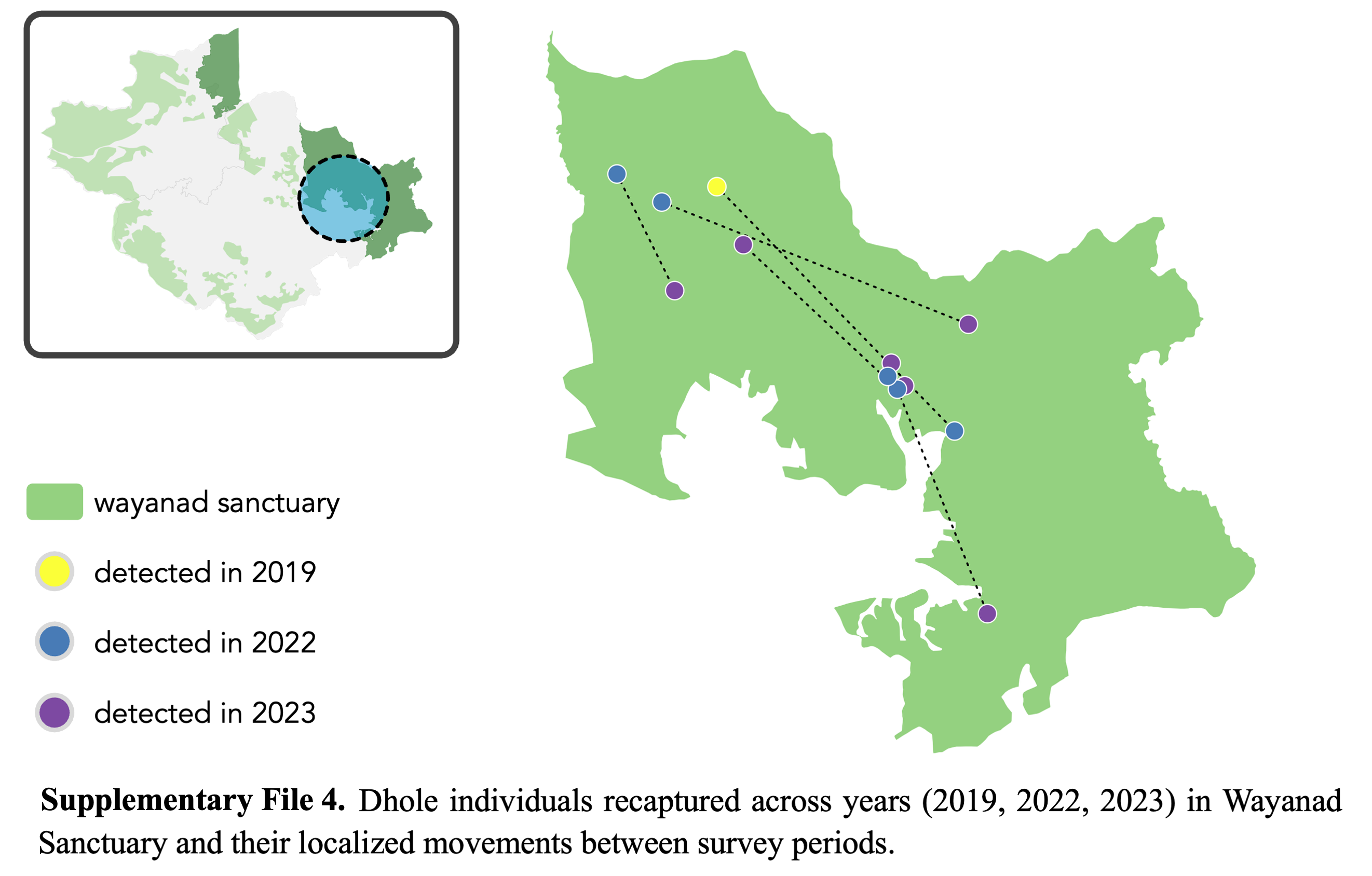
